# A Survey of Genome Editing Activity for 16 Cpf1 orthologs

**DOI:** 10.1101/134015

**Authors:** Bernd Zetsche, Jonathan Strecker, Omar O. Abudayyeh, Jonathan S. Gootenberg, David A. Scott, Feng Zhang

## Abstract

The recently discovered class 2 CRISPR-Cas endonuclease Cpf1 offers several advantages over Cas9, including the ability to process its own array and requirement for just a single RNA guide. These attributes make Cpf1 promising for many genome engineering applications. To further expand the suite of Cpf1 tools available, we tested 16 Cpf1 orthologs for activity in eukaryotic cells. Four of these new enzymes demonstrated targeted activity, one of which, from *Moraxella bovoculi* AAX11_00205 (Mb3Cpf1), exhibited robust indel formation. We also show that Mb3Cpf1 displays some tolerance for a shortened PAM (TTN versus the canonical Cpf1 PAM TTTV). The addition of these enzymes to the genome editing toolbox will further expand the utility of this powerful technology.

---

Class 2 CRISPR-Cas systems are naturally occurring microbial adaptive immune systems with single effector enzymes. The effector enzymes, such as Cas9, are RNA guided DNA endonucleases, which have been harnessed for a range of genome engineering applications (Doudna and Charpentier, 2014; Hsu et al., 2014). Although Cas9 was the first such enzyme to be developed as a genome editing tool (Cong et al., 2013; Mali et al., 2013), three orthologs of Cpf1, a single RNA-guided class 2 effector, from *Francisella novicida* U112 (FnCpf1), *Acidaminococcus sp*. BV3L6 (AsCpf1), and *Lachnospiraceae bacterium* ND2006 (LbCpf1), have also been used for genome editing in eukaryotic cells (Endo et al., 2016; Kim et al., 2016; Ma et al., 2017; Zetsche et al., 2015; Zetsche et al., 2016). Endonucleases of the Cpf1-family differ from the Cas9-family in several ways: (i) Cpf1 utilizes T-rich protospacer adjacent motifs (PAMs) located 5' of the targeted DNA sequence, (ii) target cleavage occurs distally from the PAM and results in sticky-end overhangs, (iii) Cpf1 is guided by a single CRISPR RNA (crRNA) and does not require trans-activating CRISPR RNA (tracrRNA); and (iv) Cpf1 possesses both RNase and DNase activity, which allows it to process its own CRISPR array (Fonfara et al., 2016; Zetsche et al., 2015). These features make Cpf1 particularly useful in certain situations, such as targeting AT-rich genomic regions and multiplexed gene targeting (Wang et al., 2017; Zetsche et al., 2016).

Given previous work showing that different Cas9 orthologs exhibit a range of activity in eukaryotic cells (Cong et al., 2013; Ran et al., 2015) and the potential advantages of Cpf1, we sought to identify additional Cpf1 orthologs with high activity in eukaryotic cells. Here we examine 16 new Cpf1-family proteins for nuclease activity in human cells. We identify four orthologs that can induce insertion/deletion (indel) events at targeted genomic loci. One ortholog, from *Moraxella bovoculi* AAX11_00205 (Mb3Cpf1), exhibited comparable activity to AsCpf1 and LbCpf1 when targeted sites containing TTTV (V = A, C, or G) PAMs. We also show that Mb3Cpf1 can recognize a TTN PAM, but with lower efficiency than the conserved TTTV PAM. Together, these new orthologs expand the genome editing toolbox, providing new enzymes that can be used for tailored applications.

We selected 16 uncharacterized Cpf1-family proteins with varying degrees of homology to three Cpf1 orthologs with confirmed activity in eukaryotic cells (FnCpf1, AsCPf1, and LbCpf1) (Endo et al., 2016; Zetsche et al., 2015) (Figure 1A). The direct repeat (DR) sequences of crRNAs associated with Cpf1 orthologs show high levels of homology (Figure 1B) and are predicted to fold into almost identical secondary structures (Figure 1C). The homology is particularly strong at the stem structure and the AAUU motif (Figure 1C), which is required for efficient crRNA maturation (Fonfara et al., 2016), suggesting the mechanism of crRNA maturation may be conserved within the Cpf1-family.

**Figure 1.**
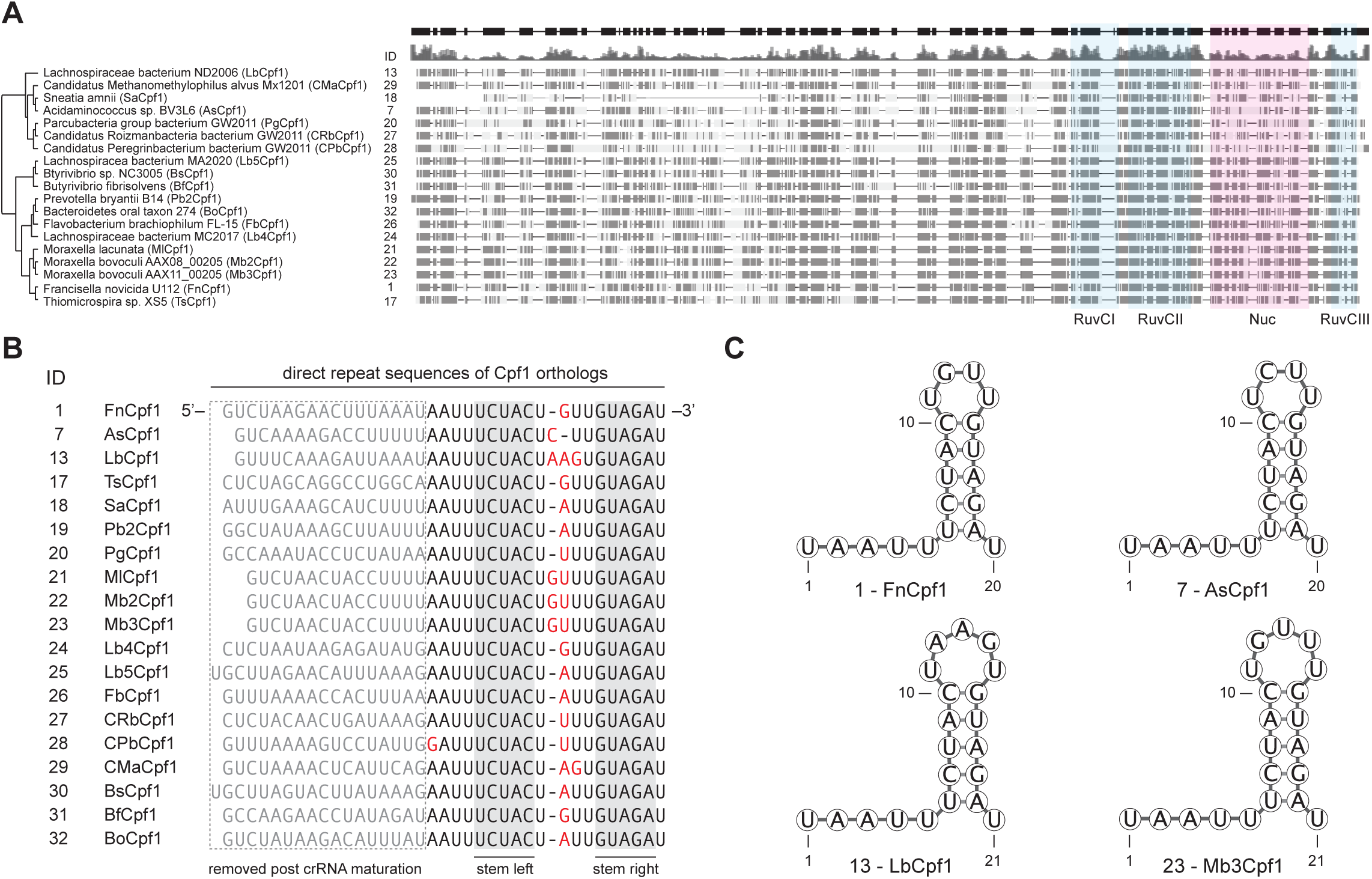
Analysis of Cpf1 ortholog diversity. A) Phylogenetic tree of 16 new Cpf1 orthologs and 3 cpf1 orthologs with confirmed activity in eukaryotic cells (1-FnCpf1, 7-AsCpf1, and 13-LbCpf1). The approximate location of the RuvC subdomains and the nuclease (Nuc) domain are shaded in blue and pink respectively. B) Alignment of direct repeat sequences of Cpf1 orthologs. Sequences that are removed post crRNA maturation are colored gray. Non-conserved bases are colored red. The stem duplex is shaded gray. C) RNA secondary structures for mature crRNAs from 1-FnCpf1, 7-AsCpf1, 13-Lbcpf1, and 23-Mb3Cpf1, predicted with the Geneious 2 software.

We performed a previously described *in vitro* assay (Gao et al., 2016) to determine the sequence of the PAM for each Cpf1 ortholog (Figure 2A). Of the 16 new Cpf1 proteins, 10 were active *in vitro* and recognized a T-rich PAM located 5' of the targeted sequence (Figure 2B), similar to previously characterized Cpf1 proteins (Zetsche et al., 2015).

**Figure 2.**
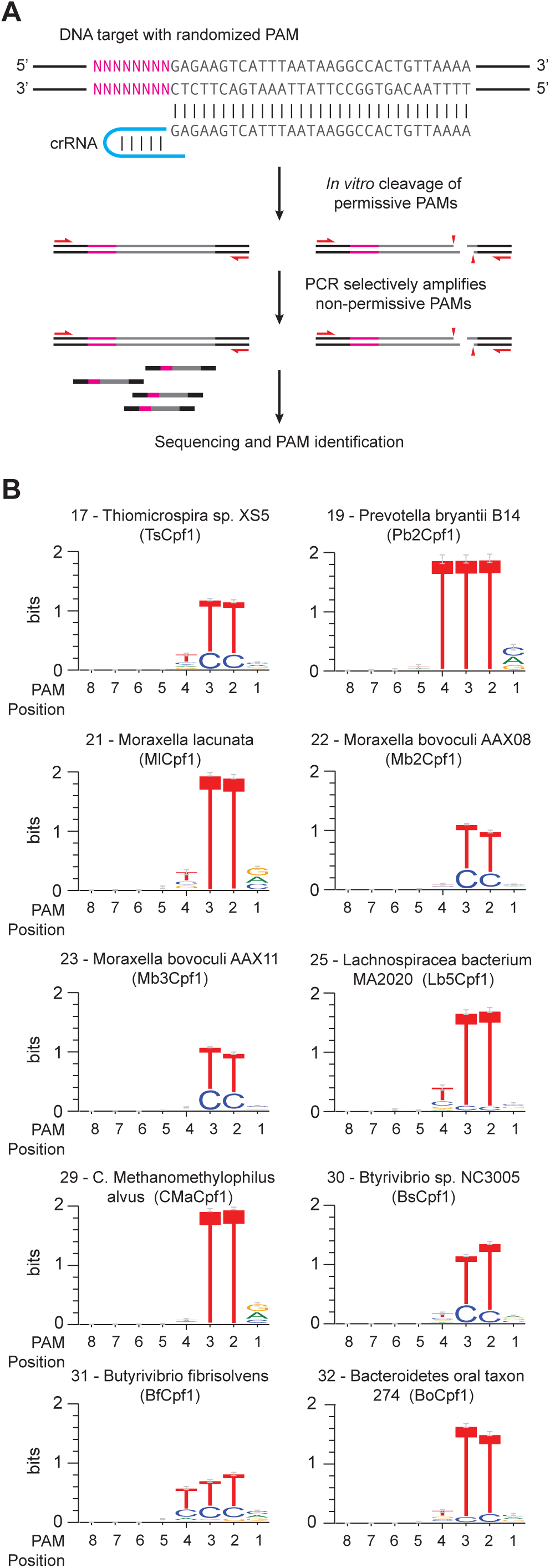
PAM identification for Cpf1 orthologs. A) Schematic for in vitro PAM screen. A library of plasmids bearing randomized 5’ PAM sequences was cleaved by individual Cpf1 nucleases and their corresponding crRNAs. Uncleaved plasmid DNA was PCR amplified and sequenced to identify depleted PAM sequences. B) PAM sequences for ten Cpf1 orthologs identified by in vitro PAM screens.

Next we tested if any of the 16 Cpf1 orthologs are active in human cells. We chose a previously validated target within *VEGFA*, located next to a TTTG PAM, permissive to all Cpf1 orthologs. HEK293T cells were transfected with plasmids encoding humanized Cpf1 orthologs together with PCR amplified fragments comprising a U6 promoter fused to the corresponding crRNA sequence (Figure 3A). Four of the new Cpf1 orthologs (*Thiomicrospira sp.* Xs5 (TsCpf1), *Moraxella bovoculi* AAX08_00205 (Mb2Cpf1), *Moraxella bovoculi* AAX11_00205 (Mb3Cpf1), and *Butyrivibrio sp*. NC3005 (BsCpf1)) were able to induce detectable indel events, as measured by surveyor nuclease assay (Figure 3B). We tested these orthologs with six additional guides targeting either *DNMT1* or *EMX1* next to TTTV PAMs (Figure 3C) and compared them to the activity of AsCpf1 and LbCpf1. For all four Cpf1 enzymes, indel frequencies of >20% could be detected for at least two guides but only Mb3Cpf1 was able to induce robust indel levels with all six guides, comparable to AsCpf1 and LbCpf1. The apparent difference in activity between Mb2Cpf1 and Mb3Cpf1 was somewhat surprising given that these orthologs share a predicted homology of 94.7%.

**Figure 3.**
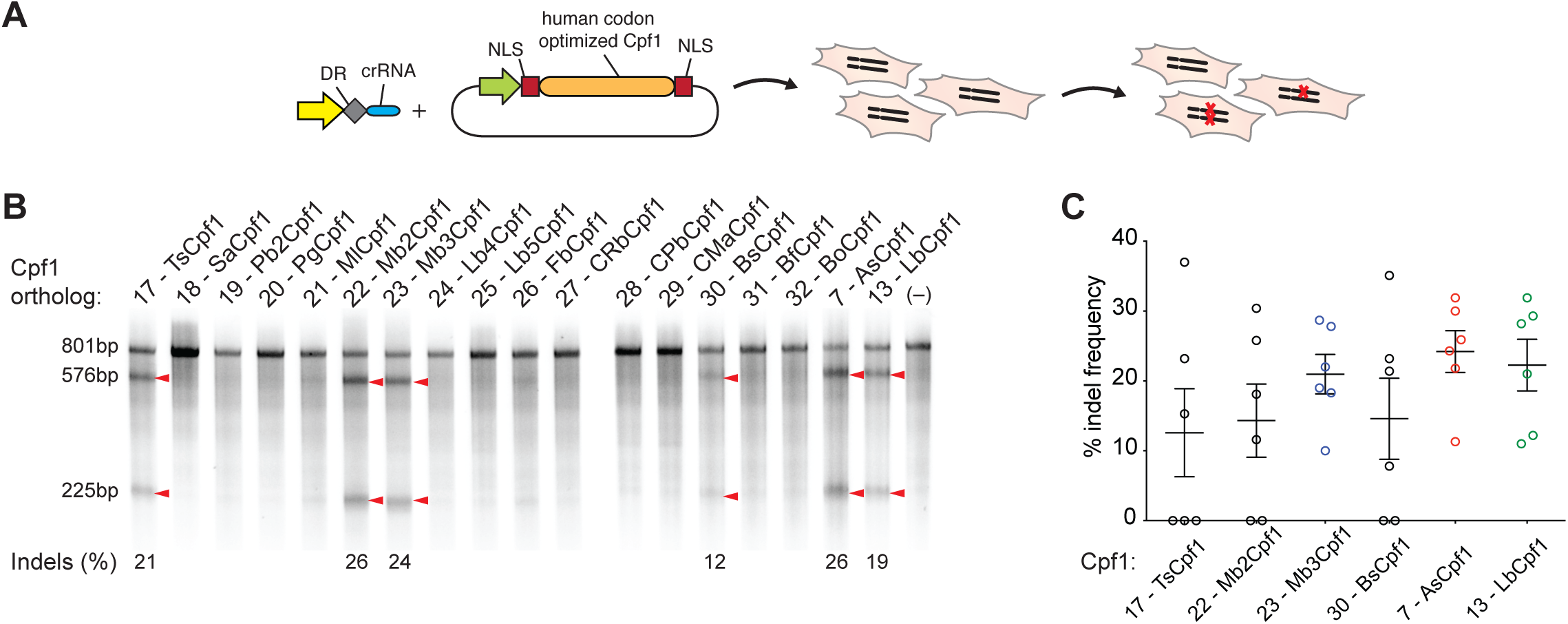
Activity of Cpf1 orthologs in human cells. A) Sixteen human codon optimized Cpf1 orthologs were expressed in HEK293T cells using CMV-driven expression vectors. The corresponding crRNA was expressed from PCR amplified fragments containing a U6 promoter fused to the crRNA sequence. B) Comparison of in vitro activity using a pre-validated guide targeting *VEGFA*, next to a TTTV (V = A, C, or G) PAM. Indel frequencies were detected by surveyor assay. Red triangles indicate cleaved fragments. Percent indel frequency is the average of three bioreplicates. C) Activity of five new Cpf1 orthologs compared to AsCpf1 and LbCpf1 using six guides targeting either *EMX1* or *DNMT1*. Each data point represents one guide; indel frequencies were determined by surveyor assay and are show as mean of all guides with SEM.

Because Mb3Cpf1 was predicted to recognize a less restrictive PAM than the TTTV consensus PAM of AsCpf1 and LbCpf1 (Figure 2B), we tested if Mb3Cpf1 can cleave endogenous DNA at TTN PAMs. To this end we designed 64 guides: 16 guides for *DNMT1*, *EMX1*, *GRIN2b*, or *VEGFA*, targeting next to any combination of NTTN PAMs. To compare the activity of Mb3Cpf1, AsCpf1, and LbCpf1 at NTTN PAMs we transfected HEK293T cells with two plasmids, one expressing Cpf1 and one expressing the crRNA, and assessed indel frequencies at each target site by deep sequencing. The average activity at TTTV PAMs was ∼18% for Mb3Cpf1, ∼28% for AsCpf1 and ∼13% for LbCpf1 (Figure 4A). A few guides targeting next to NTTN PAMs (three for MbCpf1 and one for AsCpf1) resulted in activity between 25 - 45% indels. However, while Mb3Cpf1 performed better than AsCpf1 and LbCpf1 at NTTN PAMs, the average activity was relative low with ∼5.3% for Mb3Cpf1, ∼2.7% for AsCpf1 and ∼1.4% for LbCpf1. Based on the in vitro PAM screen, Mb3Cpf1 tolerates Cs or Ts within its PAM. To assess the tolerance for Cs at position 2 and 3 of the Mb3Cpf1 PAM we used 18 guides targeting *DNMT1* or *EMX1* next to RTTN and NYYN PAMs (R = A or G, Y = C or T).

**Figure 4.**
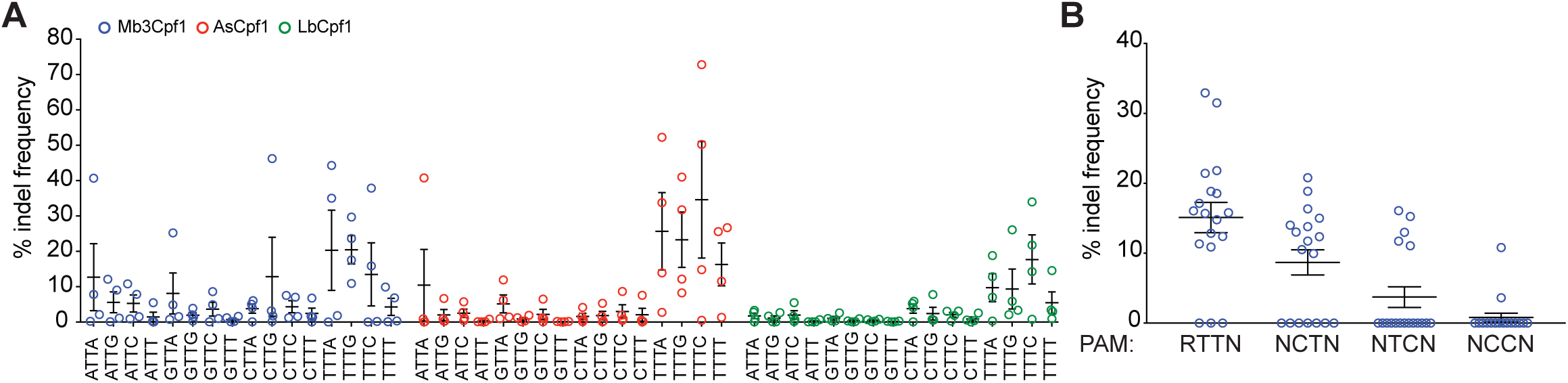
Evaluation of activity with relaxed PAM sequences. A) Mb3Cpf1, AsCpf1, and LbCpf1 were tested for recognition of NTTN PAMs using four guides per PAM, targeting four different genes (*DNMT1*, *EMX1*, *GRIN2b*, or *VEGFA*). Indel frequency was determined by deep sequencing. Each data point represents average of three bioreplicates for one guide. Data are shown as mean with SEM. B) Mb3Cpf1 was tested with 18 guides targeting either *DNMT1* or *EMX1* next to a RTTN and NYYN PAM (R = A or G, Y = C or T). Each data point represents on guide, data are shown as mean with SEM.

HEK293T cells were transfected with a single plasmid expressing Mb3Cpf1 and crRNA. The activity of each guide was determined using the surveyor nuclease assay. Guides targeting next to RTTN, RCTN and RTCN PAMs had an average activity of ∼15%, ∼9% and ∼4% respectively, while guides targeting next to RCCN PAMs are mostly inactive (Figure 4B). Taken together, our data show that Mb3Cpf1 is active in human cells and shows robust activity at TTTV PAMs, comparable to AsCpf1 and LbCpf1. Furthermore, Mb3Cpf1 can reliable target sites with RTTV PAMs albeit with lower overall activity.

Here we examined 16 new Cpf1 family proteins for potential use in genome editing. Four of these, *Thiomicrospira sp.* Xs5 (TsCpf1), *Moraxella bovoculi* AAX08_00205 (Mb2Cpf1), *Moraxella bovoculi* AAX11_00205 (Mb3Cpf1), and *Butyrivibrio sp.* NC3005 (BsCpf1), exhibited activity in human cells. Further analysis of the PAM requirements of the most active new ortholog, Mb3Cpf1, showed that it has a less restricted PAM (TTV) compared to AsCpf1 and LbCpf1, which are only active at the canonical TTTV. Previously, we observed only weak activity of FnCpf1 in mammalian cells (Zetsche et al., 2015). However, a recent study found that FnCpf1 exhibits robust activity in plant cells (Endo et al., 2016), indicating that Cpf1 orthologs might have different activity depending on the organism. The availability of another Cpf1 ortholog with high activity in eukaryotic cells will further expand the genome editing options for a wide range of organisms.

## Methods

*Computational search for Cpf1 orthologs.* Cpf1 orthologs were selected as previous described (Zetsche et al., 2015).

*Cell culture and transfection.* HEK293T cells were maintained at 37°C with 5% CO_2_ in Dulbecco Modified Eagle Medium (Gibco) supplemented with 10% fetal bovine serum (HyClone) and 2 mM GlutaMAX (Life Technology). For indel analysis 22,000 cells were seeded per 96-well (Corning) one day before transfection. Each well was transfected with 100 ng Cpf1 encoding plasmid (see Table 1) and 50 ng guide encoding plasmid or PCR fragment, or 150 ng Cpf1 and guide encoding plasmid, using Lipofectamine 2000 (Thermo Fisher Scientific). Cell were harvested 3 days after transfection with QuickExtract DNA extraction solution according to the manufacturer’s protocol and analyzed by surveyor assay or deep sequencing. For generation of Cpf1 containing whole cell lysate, 120,000 cells were seeded per 24-well (Corning) one day before transfection. Each well was transfected with 500 ng Cpf1 encoding plasmid and cell lysate was harvested 2 days after transfection.

*In vitro PAM identification assay.* The in vitro PAM identification assay was performed as described previously (Gao et al., 2016). Briefly, whole cell lysate from HEK293T cells, overexpressing one of the Cpf1 orthologs was prepared with lysis buffer (20 mM HEPES, 100 mM KCl, 5 mM MgCl_2_, 1 mM DTT, 5% glycerol, 0.1% Triton X-100) supplemented with EDTA-free cOmplete Protease Inhibitor Cocktail (Roche). CrRNA with corresponding direct repeat sequences were transcribed in vitro using custom oligonucleotides and HiScribe T7 in vitro Transcription Kit (NEB) according to the manufacturer’s recommended protocol for small RNA transcripts. The PAM library consisted of a pUC19 plasmid carrying a degenerate 8-bp sequence 5’ of a 33-bp target site (Zetsche et al., 2015). The library was pre-cleaved with XmnI and column purified prior to use (Qiagen). Each in vitro cleavage reaction consisted of 1 ul 10x CutSmart buffer (NEB), 200 ng PAM library, 500 ng in vitro transcribed crRNA, 10 ul cell lysate and water for a total volume of 20 ul. Reactions were incubated at 37°C for one hour and stopped by adding 500 ul buffer PB (Qiagen) followed by column purification. Purified DNA was amplified and sequenced using a MiSeq (Illumina) with a single-end 150-cycle kit. Sequencing results were entered into the PAM discovery pipeline (Zetsche et al., 2015).

*Surveyor assay.* Surveyor assay was performed as previous described (Hsu et al., 2013). Briefly, genomic regions flanking a target side for each gene were amplified by PCR and products were purified by QiaQuick Spin Column (Qiagen). 400 ng total purified PCR products were mixed with 2 μl 10 Taq DNA Polymerase buffer (Enzymatics) and ultrapure water to a final volume of 20 μl, and re-annealed by heating to 95°C for 2 min and slow cool down to 10°C (∼2.5°C per min). Re-annealed products were treated with surveyor nuclease (IDT) according to the manufacturer’s protocol and cleavage products were visualized on 10% Novex TBE polyacrylamide gels (Life Technologies). Gels were stained with SYBR Gold DNA stain (Life Technologies) for 10 min and imaged with a Gel Doc imaging system (Bio-Rad).

*Deep Sequencing.* Targeted regions were amplified using a previously described two-step PCR protocol (Hsu et al., 2013). Indels were counted computationally as previously described (Gao et al., 2016). Briefly, each amplicon was searched for exact matches within a 70-bp window around the cut side. For each sample, the indel rate was determined as (number of reads with indel) / (number of total reads). Samples with fewer than 1000 total reads were not included in subsequent analyses.

## Acknowledgements

We thank R. Macrae, R. Belliveau and L. Gao for discussions and support. J.S. is supported by the Human Frontier Science Program. O.A.A. is supported by a Paul and Daisy Soros Fellowship and National Defense Science and Engineering Fellowship. J.S.G. is supported by a D.O.E. Computational Science Graduate Fellowship. F.Z. is a New York Stem Cell Foundation-Robertson Investigator. F.Z. is supported by the NIH through NIMH (5DP1-MH100706 and 1R01-MH110049), NSF, Howard Hughes Medical Institute, the New York Stem Cell, Simons, Paul G. Allen Family, and Vallee Foundations; and James and Patricia Poitras, Robert Metcalfe, and David Cheng. A patent has been filed relating to the presented data. F.Z. is a founder and scientific advisor for Editas Medicine, and a scientific advisor for Horizon Discovery. Reagents will be available to the academic community through Addgene and additional information on using these reagents can be obtained via the Zhang lab website (www.genome-engineering.org).

## Author Contributions

B.Z., J.S., and F.Z. conceived this study. B.Z. and J.S. performed experiments with help from all authors. O.A. and J.G. analyzed PAM detection data. D.S. contributed to computational analysis of Cpf1 orthologs. F.Z. supervised research. B.Z. and F.Z. wrote the manuscript with input from all authors.

